# An integrative framework identifies cooperative targeting of host pathways by tick salivary miRNAs

**DOI:** 10.64898/2025.12.05.692500

**Authors:** José María Medina, Michail Kotsyfakis, Michael Hackenberg

## Abstract

Ticks are ectoparasites that modulate host responses to sustain prolonged blood feeding, and in *Ixodes ricinus*, salivary microRNAs (miRNAs) represent promising candidates for manipulating host gene expression. Using phylogenetic footprinting combined with cooperative targeting analysis, we identified deeply conserved miRNA–mRNA interactions that appear essential for the tick lifecycle and its ability to parasitize diverse vertebrate hosts. Our findings show that conserved tick miRNAs can mimic host miRNAs by exploiting shared target sites on host transcripts, allowing them to modulate regulatory circuits critical during tick feeding. We identified twelve core salivary miRNAs—highly expressed and enriched in conserved target sites—that cooperatively target twenty-two human genes predominantly expressed in skin and neural tissues. This cooperative suppression converges on host hub genes like *PDGFRA* and *NRG1*, linking tick miRNA activity to MAPK and PI3K–AKT pathways that regulate immune defense, tissue repair, and sensory responses. Overall, our results demonstrate that *I. ricinus* miRNAs can synergistically disrupt host homeostasis, and they introduce a broadly applicable framework that leverages evolutionary conservation to detect meaningful cross-species miRNA–mRNA interactions.

**Highlights:** The study shows that tick miRNAs with Bilaterian node of origin mimic host miRNAs to exploit conserved target sites of host mRNAs.

It demonstrates that these conserved tick miRNAs cooperatively repress key host defense and repair pathways.

It introduces a framework that leverages evolutionary conservation to detect and explain functional, conserved cross-species miRNA–mRNA regulation.

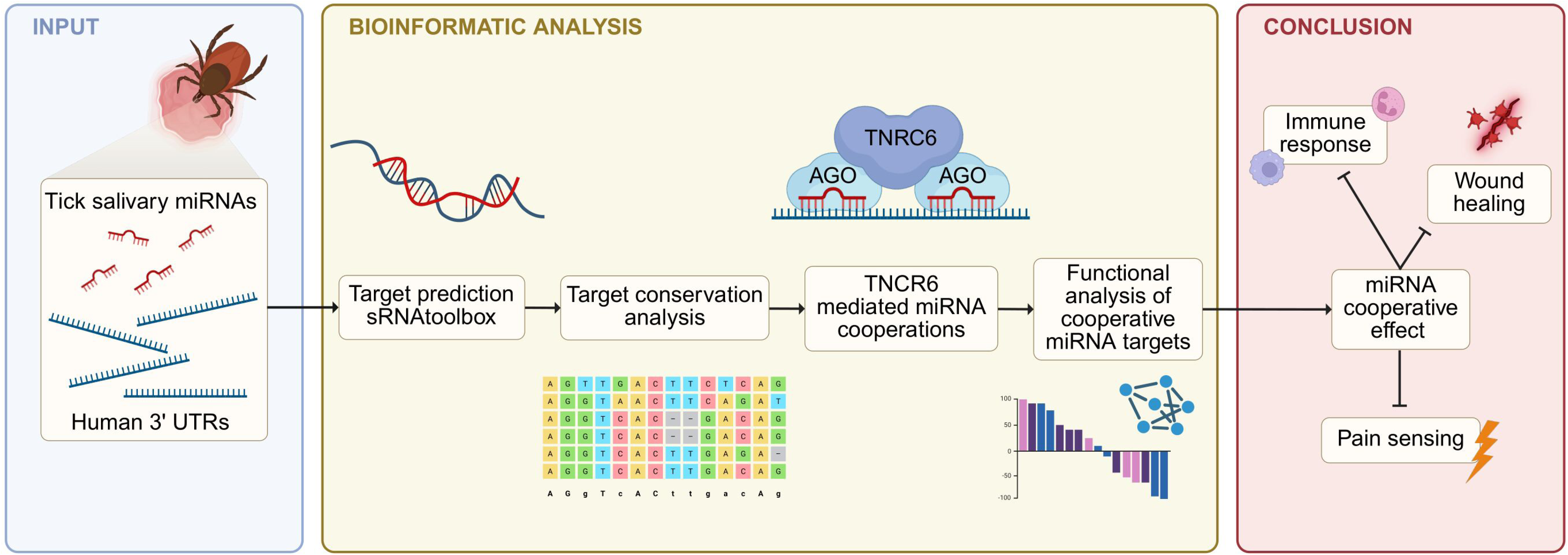

## Introduction

*Ixodes ricinus*, commonly known as the castor bean tick, is a prominent ectoparasite with significant implications for both animal and human health (Sprong et al., 2018). This tick species, similar to many other tick species, is known for its broad range of hosts, including mammals, birds, and reptiles (Gern & Humair, 2002), which contributes to its wide geographical distribution across Europe and parts of North Africa (Gern & Humair, 2002; Zhioua et al., 1999). Recently, the prevalence of *I. ricinus* and tick-borne diseases in humans has been increasing, a trend that the research community attributes to environmental factors such as climate change, over-urbanization, and the alteration of the natural habitats of ticks (Gilbert, 2021; Madison-Antenucci et al., 2020). These factors enhance tick survival and facilitate their expansion into new green-urban areas (Hansford et al., 2022; Medlock et al., 2013). Therefore, further research on the parasitic mechanisms mediating tick feeding success is of critical importance.

Ticks penetrate host skin to feed, which triggers wound healing and immune responses in the host (Hajnická et al., 2011; Kahl & Gray, 2023; Kotál et al., 2015). To counter these host defenses and ensure blood flow, ticks inject saliva with immunomodulatory and anti-hemostatic molecules while drawing blood (Francischetti et al., 2009; Šimo et al., 2017). Extensive research has been conducted to identify and characterize components of tick saliva that modulate host response, with a particular focus on tick salivary proteins and tick salivary gland transcriptomics (Abbas et al., 2022; Chmelar et al., 2012; Chmelař et al., 2016; Kotsyfakis et al., 2015; Medina, Abbas, et al., 2022; Medina, Jmel, et al., 2022). However, among the diverse tick salivary components are also microRNAs (miRNAs) (Hackenberg et al., 2017; Nawaz, Malik, Zhang, Gebremedhin, et al., 2020), small non-coding RNAs that post-transcriptionally regulate gene expression (Bartel, 2004). These molecules have emerged in the last decade as molecular effectors of cross-species and even cross-kingdom communication across biological systems (Arcà et al., 2019; Buck et al., 2014; Cano-Argüelles et al., 2023; Chowdhury et al., 2024; Hackenberg et al., 2017; M. Wang et al., 2016). For instance, helminths can secrete exosome-packaged small RNAs that enter mammalian cells and suppress innate immunity (Buck et al., 2014). Plants are also able to engage in bidirectional small RNA exchange with fungal pathogens, using this mechanism as a form of defense (M. Wang et al., 2016). In hematophagous arthropods, saliva-derived mosquito miRNAs have been shown to mimic host miRNAs and may modulate host–pathogen interactions (Arcà et al., 2019). Likewise, ticks salivary miRNAs are predicted to target host genes involved in immunity and inflammation, potentially influencing host responses at the bite site (Cano-Argüelles et al., 2023; Hackenberg et al., 2017).

Despite their potential as gene expression regulators, accurate prediction of miRNA-mRNA interactions remains a challenge. Most of the studies described above have inferred miRNA–mRNA interactions using prediction algorithms that depend on partial complementarity in the seed region, but such approaches often overestimate miRNA targets and are associated with high rates of false positives (Loganantharaj & Randall, 2017; Pinzón et al., 2017). Recent computational approaches aim to improve accuracy by integrating evolutionary conservation, mRNA accessibility, and the clustering of binding sites regions (Agarwal et al., 2015; Aparicio-Puerta et al., 2022). However, these methods are often optimized for model organisms and have not been broadly applied to parasitic systems (Agarwal et al., 2015).

Beyond individual targeting, growing evidence suggests that miRNAs may act cooperatively to enhance gene regulation (Briskin et al., 2020; Diener et al., 2023; Kilikevicius et al., 2021; Sætrom et al., 2007). Prior analysis of *I. ricinus* salivary miRNAs revealed co-targeting of specific genes more often than expected by chance (Hackenberg et al., 2017). Given the stochastic nature of miRNA-mediated repression, cooperative action—where multiple miRNAs target the same transcript—can amplify regulatory effects (Kilikevicius et al., 2021). This phenomenon has a molecular basis: AGO–miRNA complexes can bind simultaneously to the scaffolding protein TNRC6, which stabilizes interactions at closely spaced target sites and enhances silencing (Briskin et al., 2020).

In this study, we present a pipeline based on the hypothesis that *I. ricinus* miRNAs parasitize conserved regions in a broad range of host species. Additionally, the workflow includes a framework to assess cooperative miRNA targeting by analyzing shared, conserved and clustered binding sites across conserved 3′ UTRs. Our findings support a model in which tick miRNAs leverage both conserved and cooperative interactions to modulate key host pathways involved in immunity, wound healing, and sensory perception, as a strategy to enhance parasitic success.

## Results

### *Ixodes ricinus* miRNA target sites in the human transcriptome

In order to detect and analyze potential interactions between microRNAs (miRNAs) from *Ixodes ricinus* and mRNAs from *Homo sapiens*, we first generated a consensus based *in-silico* target prediction. A total of 380,696 target sites in the UTRs of *H. sapiens* were predicted for the set of 162 mature *I. ricinus* miRNAs. As a negative control, we constructed a set of miRNAs with randomized sequences which preserve the base composition but were no allowed to have known seed regions in MirGeneDB (Clarke et al., 2025). The randomized miRNAs yielded a total of 353,239 predicted target sites in the UTRs of *H. sapiens*.

The difference between the predicted target sites of *I. ricinus* miRNAs and the randomized set of miRNAs is illustrated in Figure 1A. Overall, *I. ricinus* miRNAs tend to have a higher number of predicted target sites compared to the randomized miRNAs, especially in the middle range of the distribution. However, it is remarkable that randomized miRNAs can occasionally achieve an exceptionally high number of predicted target sites purely by chance, even exceeding those observed in *I. ricinus* miRNAs. This confirms that target prediction can produce a high number of false positives.

**Figure 1.**
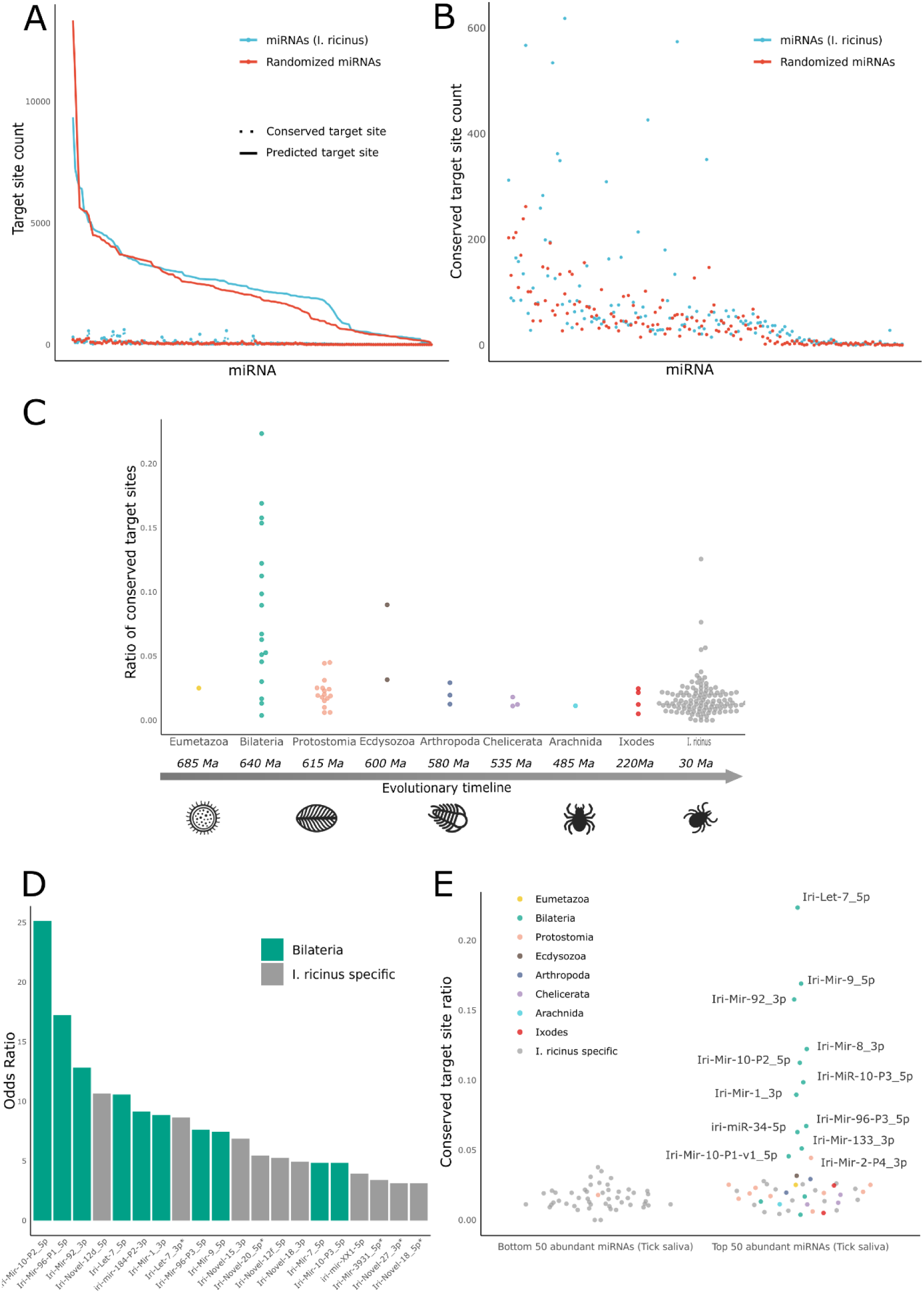
Prediction and conservation analysis of Ixodes ricinus microRNAs target sites in H. sapiens 3’ UTRs. **A)** Relationship between predicted and conserved microRNA target sites in I. ricinus and randomized miRNAs. The X-axis represents pairs of miRNAs, each consisting of one I. ricinus miRNA and one randomized miRNA, ordered by their number of predicted target sites. The Y-axis shows the count of predicted (line) and conserved (dots) target sites, respectively. The blue line represents I. ricinus miRNAs, while the red line corresponds to the randomized miRNAs. **B)** Comparison between conserved target sites of I. ricinus miRNAs and the control set of randomized miRNAs, respectively. This figure presents a magnified view of the dot plot shown in Figure 1A. The Y-axis represents the number of conserved target sites, and the X-axis shows pairs of miRNAs ordered by their number of predicted target sites. **C)** Evolutionary conservation of I. ricinus microRNAs and their conserved-to-predicted target ratios. The Y-axis represents the ratio of conserved mRNA target sites to the total predicted target sites. The X-axis represents the taxonomic level of seed sequence conservation, with deeper taxa positioned to the left. An evolutionary timeline is shown to provide a reference for the divergence times of the taxa mentioned, expressed in millions of years. **D)** Top 20 microRNAs showing greater target conservation than randomized expectation. The Y-axis represents the odds ratio obtained from Fisher’s exact test, comparing I. ricinus miRNAs with randomized miRNAs after ranking both by the number of predicted target sites. The taxonomic node of origin of the miRNA is indicated by different colors. **E)** Ratio of conserved target sites between the least and most abundant microRNAs in I. ricinus saliva. The 50 most abundant and the 50 least abundant miRNAs in tick saliva are represented in this plot. The Y-axis shows the ratio of conserved mRNA target sites to the total predicted target sites. The nodes of origin are indicated by different colors.

To reduce the number of false positives, we filtered the predicted targets by a phylogenetic footprinting approach. This analysis was motivated by the fact that *I. ricinus* has a wide range of diverse potential hosts, including humans, large and small mammals, birds, and reptiles (Gern & Humair, 2002). Consequently, if tick miRNAs play a role in the feeding success on a broad range of hosts, the target sites need to be conserved.

From the 380,696 predicted target sites, 11,746 (3.08%) were identified as conserved. In contrast, only 7,454 conserved target sites (2.11%) were identified from the 353,239 predicted target sites for the randomized miRNAs. The proportion of conserved target sites for *I. ricinus* miRNAs was significantly higher than that for the randomized set (p < 0.00001). This significant difference suggests a critical importance of the phylogenetic footprinting analysis in this study.

Figure 1A and its magnified view in Figure 1B illustrate that miRNAs with the highest number of target sites do not necessarily exhibit the highest number of conserved target sites. This finding hints towards the existence of some mechanism and against the spurious random detection of conserved targets. Notably, high ratios were observed only for *I. ricinus* miRNAs and similar values do not appear in the distribution of the randomized set of miRNAs.

### Evolutionary conservation of tick miRNAs and saliva abundance

After the knock-out of highly conserved miRNAs, severe phenotypes like embryonic lethality and different developmental defects among many others can be observed(Bartel, 2018). This shows that deeply conserved miRNAs are generally more important than evolutionary younger ones. Furthermore, it can be expected that functionally relevant tick miRNAs should be secreted during tick feeding and therefore present in saliva.

To test whether higher ratios of conserved target sites might indicate functional relevance, we analyzed this property as a function of conservation depth, i.e. node of origin and saliva abundance. We assigned 22 mature tick microRNA sequences to the node of Bilateria, which are therefore shared across all bilaterian animals and present in host species, typically having between 0 and 3 evolutionary changes outside the seed region. The rest of the mature tick sequences are shared by all Protostomates (17), Ecdysozoa (3), Arthropoda (3), Arachnida (6), Chelicerata (13) or seem to be tick specific (see Supplementary File 1). Note that many guide sequences of deeply conserved microRNAs are labeled as tick specific because they have seed sequences not observed in any other microRNAs in MirGeneDB.

Figure 1C displays that deeply conserved tick miRNAs, i.e. those shared across bilaterian animals tend to have higher ratios of conserved targets which suggests a relation between target conservation and functional importance. Interestingly, a ranked comparison to determine which miRNAs have more conserved target sites than expected by chance identified two statistically significant groups (Figure 1D, Supplementary File 2). The first group comprises bilaterian miRNAs, while the second group consists of miRNAs annotated only in *I. ricinus* which have emerged more recently or even might be restricted to *I. ricinus*.

To further analyze whether the ratio of conserved target sites is indicative of its functional importance in the tick-host interaction, we examined if this ratio correlates with miRNA abundance in tick saliva (Hackenberg et al., 2017). The comparison between the 50 most abundant miRNAs in saliva and the 50 least abundant miRNAs, as shown in Figure 1E, revealed that several miRNAs highly abundant in tick saliva exhibit a significantly higher ratio of conserved target sites compared to lesser abundant in tick saliva. Additionally, those miRNAs with the highest ratio of conserved target sites were found to be conserved in bilaterian animals.

### In silico functional analysis of host target genes

To establish a baseline for the functional impact of tick miRNAs on host parasitism, we first asked whether functional annotations enriched in target genes could be detected when considering all *I. ricinus* miRNAs. A total of 20061 genes had at least one target site identified for at least one of the 162 *I. ricinus* miRNAs while this number drops to 4938 when only considering conserved targets. Additionally, to focus on those genes with the highest potential to be modulated by a specific miRNA we analyzed a set of 526 genes which had 5 identified conserved target sites or more. These genes were functionally characterized, and their pathway and ontology enrichment were analyzed by pathway enrichment analysis using the KEGG, Reactome, GO and WikiPathways databases. The analysis revealed that this set of genes was significantly enriched in 16 KEGG pathways, 45 Reactome pathways, 351 GO terms from the category Biological process, 41 from the category Molecular Function and 112 from the category Cellular Component (Supplementary Figure 1B) and 35 WikiPathways pathways. The high number of pathways with statistically significant enrichment suggests a microRNA-based strategy to manipulate multiple host homeostatic responses to a feeding tick (Supplementary Figure 1A).

### Cooperative effect analysis

It has been demonstrated that miRNAs may act synergistically in gene expression regulation (Kilikevicius et al., 2021), that, in the case of ticks, can lead to a cooperative effect in host homeostasis modification. This synergy can enhance repression strength, allowing therefore that even a limited number of miRNAs exert a significant influence on their target mRNAs, which may be particularly relevant for *I. ricinus*, where delivery to host cells may be quantitatively limited (Chevillet et al., 2014; Iftikhar & Carney, 2016; Turchinovich et al., 2016). Therefore, we examined the potential cooperative effects of the complete set of *I. ricinus* miRNAs, focusing on their ability to target conserved neighboring sites within the same host mRNA. Such a close arrangement of conserved target sites facilitates the simultaneous attachment of AGO-miRNA complexes to the TNRC6 protein. Our analysis demonstrates that 755 target sites (6.43%) out of 11,746 conserved target sites were identified as cooperative, forming 451 distinct synergistic interactions. In comparison, a randomized miRNA set showed that 394 of 7,454 conserved target sites (5.29%) could cooperate, forming 218 unique interactions. The difference in cooperative target regions between *I. ricinus* miRNAs and randomized miRNAs is significant, as determined by Fisher’s exact test (p = 0.0003).

Next, we examined the distribution of cooperative target sites relative to conserved target sites across *I. ricinus* miRNAs and the randomized (control) miRNAs (Figure 2A, B). Unlike the pattern observed for conserved and total target sites, miRNAs with more conserved target sites tend to also have more cooperative target sites. This is true for both, tick miRNAs and the control set, having however the set of *I. ricinus* miRNAs a higher number of cooperative target sites. Regarding the ratio of cooperative to conserved target sites per specific miRNA, no significant differences were observed between the *I. ricinus* and randomized miRNA sets. Notably, miRNAs with the highest ratios of cooperative target sites were *I. ricinus*-specific (Figure 2C), with only one *I. ricinus* miRNA conserved in protostomes, namely Iri-Mir-2-P4_3p, showing more cooperative target sites than expected by chance (FDR: 0.032, Supplementary File 3).

**Figure 2.**
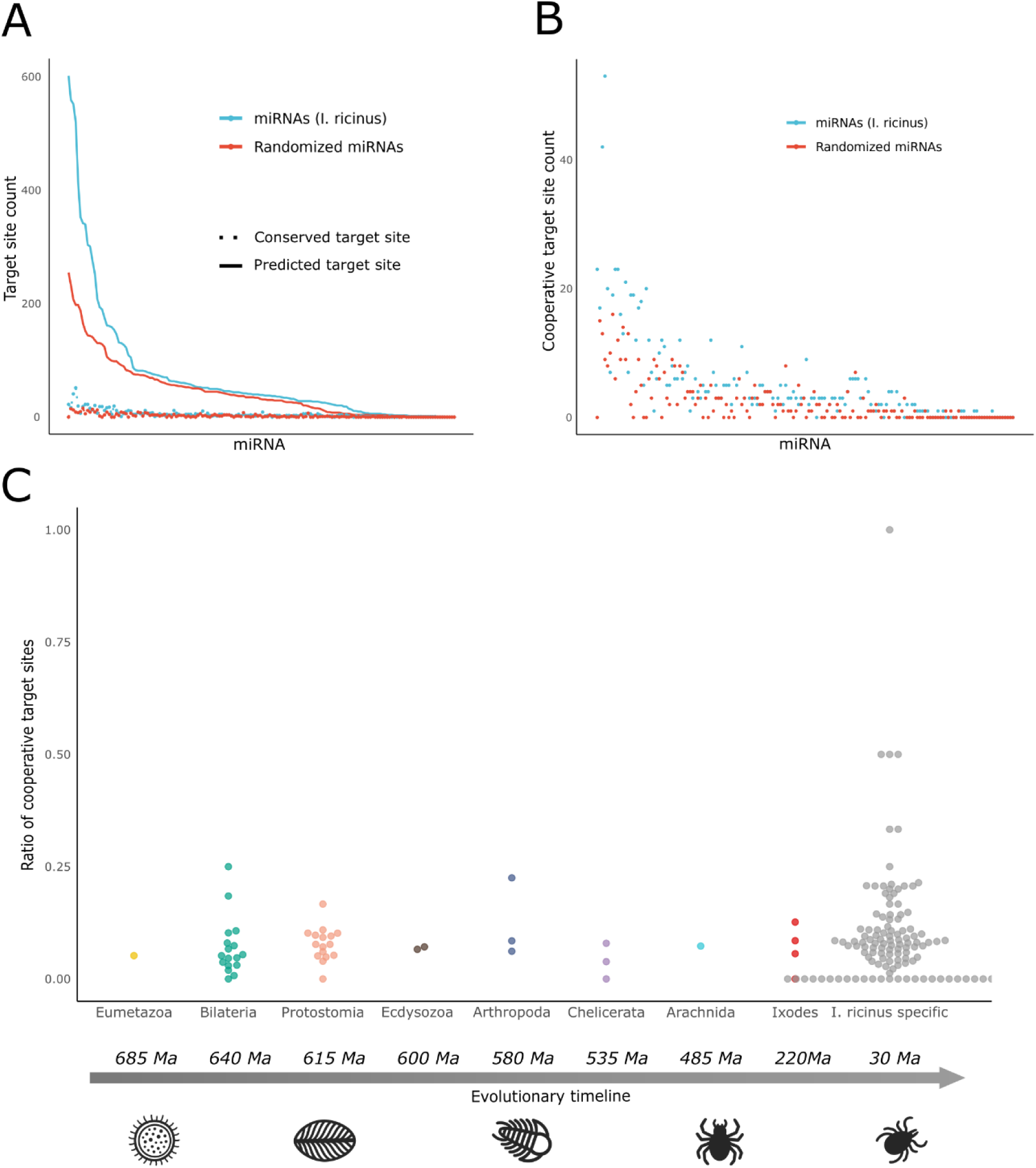
Analysis of cooperation between microRNAs of Ixodes ricinus. **A)** Relationship between conserved and cooperative miRNA target sites in I. ricinus and randomized miRNAs. The X-axis represents pairs of miRNAs, each consisting of one I. ricinus miRNA and one randomized miRNA, ordered by their number of conserved target sites. The Y-axis shows the count of conserved and cooperative target sites. The blue line represents I. ricinus miRNAs, while the red line corresponds to the randomized miRNAs. **B)** Comparison between cooperative target sites of I. ricinus miRNAs and the control set of randomized miRNAs, respectively. This figure presents a magnified view of the dot plot shown in Figure 2A. The Y-axis represents the number of cooperative target sites, and the X-axis shows pairs of miRNAs ordered by their number of conserved target sites. **C)** Node of origin of I. ricinus miRNAs and their cooperative-to-conserved target ratios. The Y-axis represents the ratio of cooperative mRNA target sites to the conserved target sites. The X-axis represents the microRNAs grouped into different taxonomic levels regarding their node of origin. An evolutionary timeline is shown to provide a reference for the divergence times of the taxa mentioned, expressed in millions of years.

### Identification of salivary miRNAs which target conserved 3’UTR sites

We next examined whether the most abundant tick salivary miRNAs (as described in our previous work (Hackenberg et al., 2017)) that also exhibit a high ratio of conserved potential target sites regulate conserved host gene targets in a synergistic manner. Thus, we identified 12 *I. ricinus* miRNAs, hereafter named as core salivary miRNAs, that stand out a) in their abundance in tick saliva, b) in their high ratio and statistical significance regarding the number of conserved target sites, c) in targeting more conserved target sites than expected by chance (Supplementary File 2) (Figure 1E). These core *I. ricinus* salivary miRNAs—namely Iri-Let-7_5p, Iri-Mir-1_3p, Iri-Mir-2-P4_3p, Iri-Mir-8_3p, Iri-Mir-9_5p, Iri-Mir-10-P1-v1_5p, Iri-Mir-10-P2_5p, Iri-Mir-10-P3_5p, Iri-Mir-34_5p, Iri-Mir-92_3p, Iri-Mir-96-P3_5p, and Iri-Mir-133_3p—are, with the exception of Iri-Mir-2-P4_3p (conserved only at the Protostomia taxon level), conserved in all bilaterian animals. Six of them shared identical sequences with their *H. sapiens* homologous miRNAs, whereas the remaining ones showed between two and three nucleotide differences in the non-seed regions (Supplementary File 4). All of them were confirmed to be present in the most recent *I. ricinus* genome assembly (Cerqueira de Araujo et al., 2025).

In total, these twelve miRNAs had 38952 predicted target sites, from which a total of 3830 were identified as conserved (9.8%). These conserved target sites were distributed among 2662 different genes. The number of genes can influence the p-values obtained in a functional enrichment analysis. Therefore, we analyzed the 526 genes with the highest number of conserved target regions for the 12 specific miRNAs in order to adjust the number of genes to the one from the previous analysis (Figure 2).

Pathway enrichment analysis revealed that these genes were associated with 10 KEGG pathways, 17 Reactome pathways, and 16 pathways from the WikiPathways database. Ontology enrichment analysis identified 307 enriched GO terms in the Biological Process category, 32 in the Molecular Function category, and 8 in the Cellular Component category. Most significant pathways and ontology terms are shown in Supplementary Figure 2A and Supplementary Figure 2B, respectively.

These results raise the question to what extend those 12 selected core salivary miRNAs are responsible for the predicted functions of the whole *I. ricinus* miRNA panel. To address this, we compared the results to those obtained using the full set of *I. ricinus* miRNAs (Supplementary Figure 2C,D). Overall, 31.5% of the pathways enriched in genes targeted by the full set of *I. ricinus* miRNAs are also enriched in the subset of genes targeted by the 12 core salivary miRNAs. Notably, when looking at database level, 62.5% of the KEGG pathways identified analyzing the function of the full set of miRNAs are covered by the core salivary miRNAs (Supplementary Figure 2C). Regarding the ontology analysis, 49.9% of the GO terms identified from the full set of miRNA targets are retained in the enrichment profile of the 12 core salivary miRNAs (Supplementary Figure 2D). These results confirm the importance of these 12 salivary miRNAs.

### Cooperative effect analysis of core *Ixodes ricinus* salivary miRNAs

Having provided evidence for the importance of these 12 miRNAs in tick-host interaction the next question is to address it capacity to act cooperatively on the target genes, thus possibly enhancing the repression strength. We identified 45 cooperative target sites located in 22 different host genes. Of these, 10 genes were associated with innate immunity, including pathways that promote inflammation and immune cell activity such as macrophages, T cells, and Natural Killer cells (Arodz et al., 2013; Dayam et al., 2017; Erdoğan et al., 2016; Forwood et al., 2007; Hayashi et al., 2015; Hock et al., 2014; Kusama et al., 2025; Qin et al., 2024; Ye et al., 2020). 5 host genes were implicated in wound healing and angiogenesis, processes that could facilitate wound closure and limit tick feeding (Kataria et al., 2019; Ki et al., 2020; Veugelers et al., 1999; X. Wang et al., 2020; Yao et al., 2022). Another 5 of these host genes were linked to pain perception and mechanosensation, both of which are critical host responses to prolonged tick attachment and feeding (Jin et al., 2023; Kádková et al., 2024; Lee et al., 2013; Liu et al., 2020). A description of the genes, their functions, and supporting references is provided in Supplementary File 5.

If the detected interactions are indeed functionally relevant, then the targeted genes must be expressed in tissue types that come into contact with the tick. We observe that the cooperative core miRNA target set is broadly expressed in skin tissue cells (Figure 3A). Targeted host genes such as SRSF10, CELF1, RIC1, GPC6, and NFIA were abundant across all skin cell types, except mast cells. Of note, mast cells are known targets for tick salivary proteins (Kotál et al., 2015). Certain cell types, such as fibroblasts, which are key players in wound healing and pro-inflammatory response (Bainbridge, 2013; Schuster et al., 2021), expressed nearly all the targeted genes, with only one gene apparently not expressed in fibroblasts. Analysis of GTEx data revealed that 12 out of 16 skin cell types, at the single cell level expression analysis, including adipocytes, lymphatic and vascular endothelial cells, basal and suprabasal keratinocytes, fibroblasts, dendritic cells, macrophages, Langerhans cells, melanocytes, pericytes, and sebaceous and sweat gland cells, exhibited a significantly higher proportion of single cells expressing these host genes than what could be expected by chance in random sets of 22 genes (Figure 3A). The same figure demonstrates that lymphatic endothelial cells and sebaceous and sweat gland cells showed significantly high expression levels of these genes. These findings add evidence one the importance of the 12 core *I. ricinus* salivary miRNAs in the host’s skin homeostasis.

**Figure 3.**
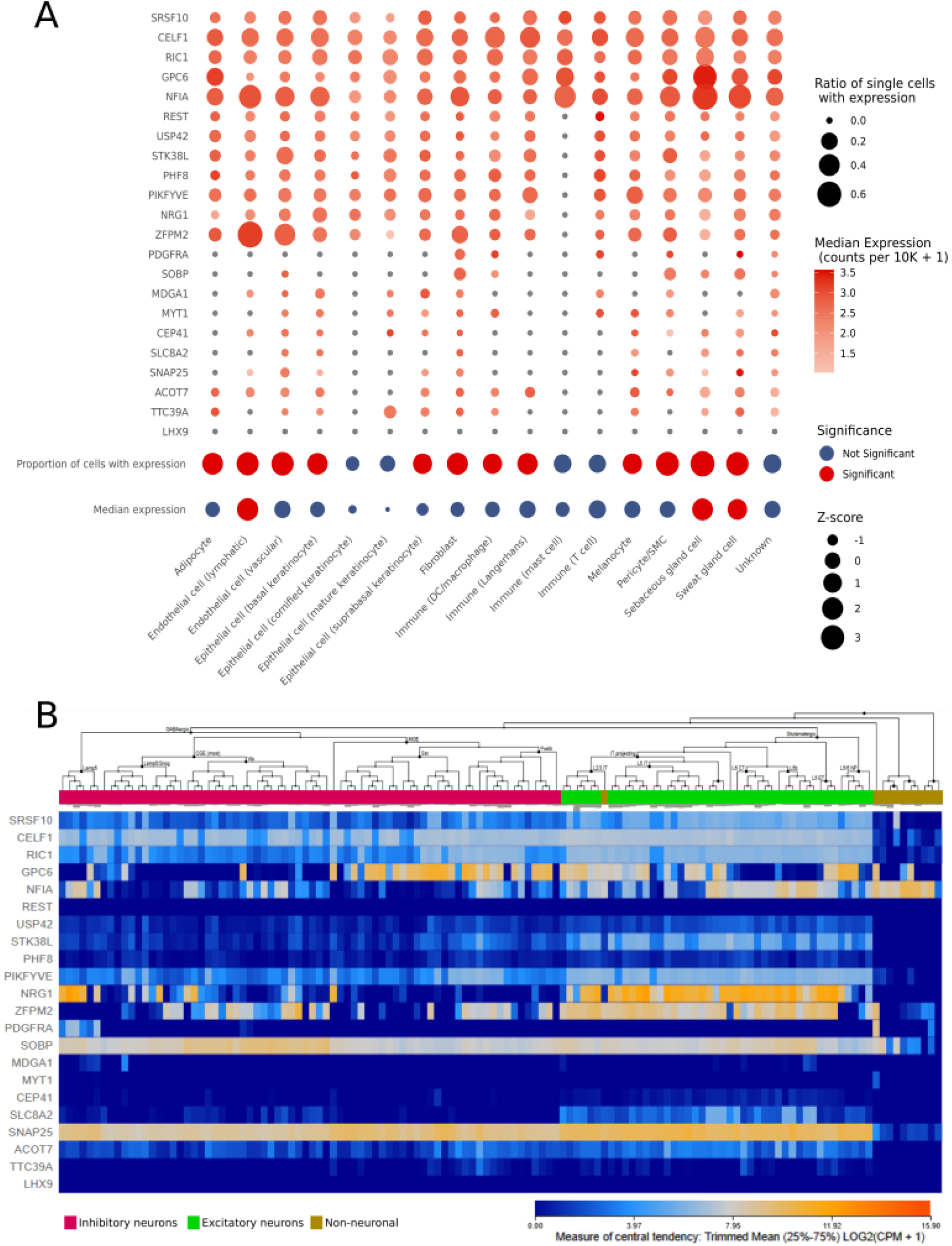
Abundance in human cells of mRNAs with conserved and cooperative target sites for core Ixodes ricinus salivary microRNAs. **A)** Abundance patterns and statistical significance in single cell types from skin tissue. The analysis was performed using the GTEx single-cell dataset (GTEx Consortium, 2013). The top part of the figure shows the abundance of genes (rows) across single-cell types (columns), where bubble size represents the proportion of single cells within each cell type expressing a given gene, and color intensity indicates median expression (counts per 10K). The bottom part illustrates the statistical significance of the median expression and the median proportion of cells expressing the genes, with red circles indicating statistically significant enrichment. Z-scores are represented by bubble size. **B)** mRNA abundance in neural cells. Inhibitory neurons, excitatory neurons and non-neuronal cells are highlighted with different colors. A color gradient between blue (low) and orange (high) represents the average mRNA expression value, measured as the log-transformed counts per million (log2(CPM + 1)), after removing the bottom 25% and top 25% of values across genes or samples.

Additionally, it would certainly be advantageous for the tick to impair the activity of sensory neurons, since they mediate the detection of an infesting tick. To investigate this possibility, we examined the expression of the 22 target genes in neural cells and found that genes cooperatively targeted by the 12 *I. ricinus* miRNAs are indeed expressed in these cells (Figure 3B). This supports the idea that tick salivary miRNAs can modulate neural cells and, consequently, may affect the function of sensory neurons in the bite region.

Based on our findings that the targeted host genes are expressed in tissues relevant to tick-host interaction, we asked whether they are functionally connected—i.e., whether they participate in the same pathways or interact at the protein level—such that their coordinated repression could synergistically affect specific host defense mechanisms and processes. We conducted a protein-protein interaction (PPI) network analysis based on the 22 target genes (Figure 4) adding 50 intermediate interactions to detect indirect relationships between these genes. PDGFRA and NRG1 showed the highest eigenvector centrality and the greatest number of interactions with other genes—24 and 15 interactions, respectively—with eigenvector centrality values of 0.269 and 0.181. In comparison, the mean number of interactions per node was 8.25, and the mean eigenvector centrality across the network was 0.073, underscoring the central roles of PDGFRA and NRG1 in this PPI network.

**Figure 4.**
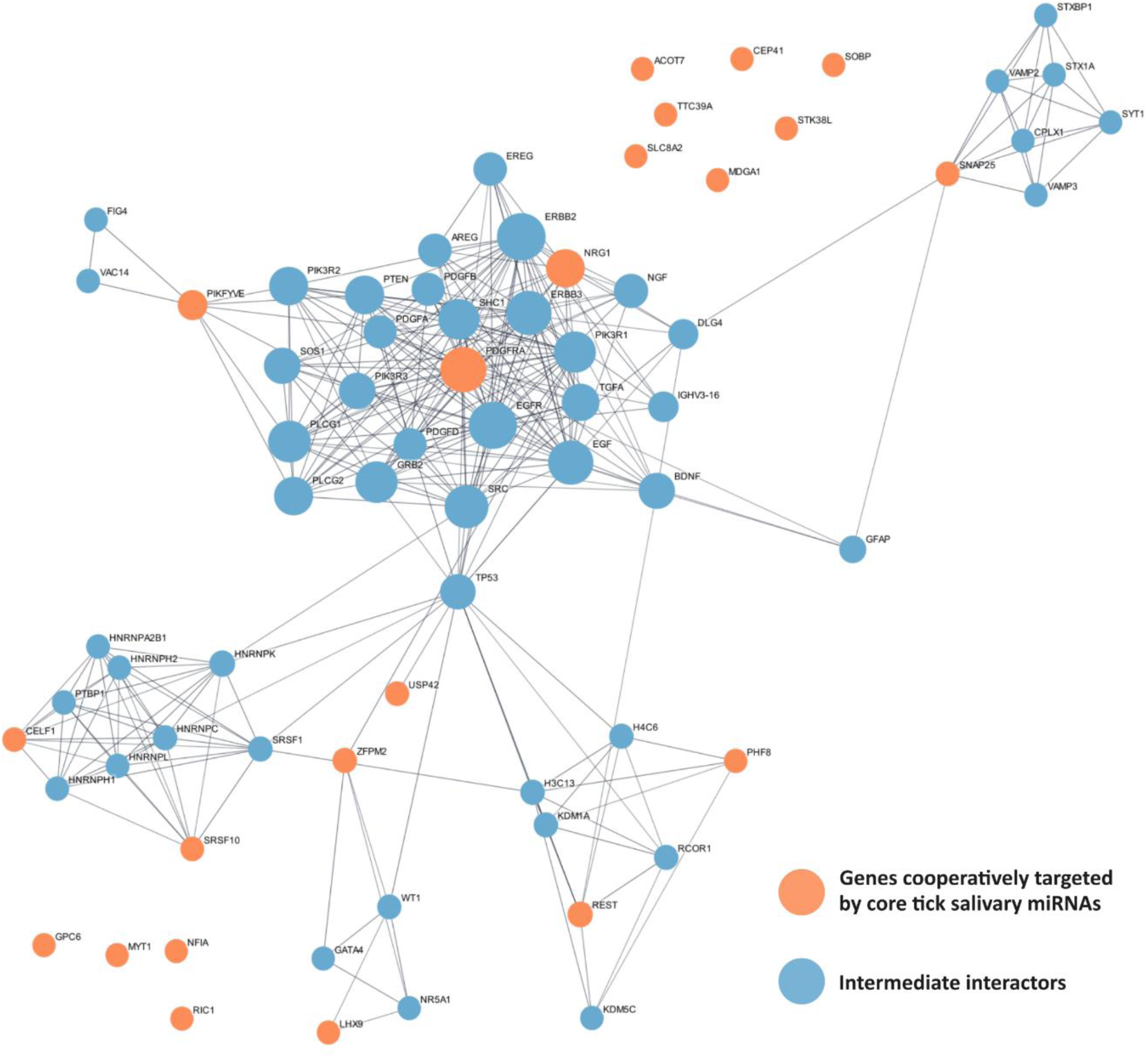
Protein-protein interaction network of gene products ‘conservedly’ and cooperatively targeted by core Ixodes ricinus salivary microRNAs. The set of genes ‘conservedly’ and cooperatively targeted by core I. ricinus salivary miRNAs are represented by color orange. Fifty interacting proteins were added using StringDB with a high-confidence threshold of 0.7 and are shown in blue.

### Hub genes involved in the cooperative effect of *Ixodes ricinus* salivary microRNAs

As NRG1 and PDGFRA are cooperatively targeted by tick miRNAs and are central in the PPI network we further focused our analysis on the involved tick miRNAs, their conservation and targeting patterns.

NRG1 is targeted at conserved regions by four mature tick miRNAs: Iri-Mir-96-P3_5p, Iri-Mir-8_3p, Iri-Mir-92_3p, and Iri-Mir-10-P3_5p. The first two, Iri-Mir-96-P3_5p and Iri-Mir-8_3p, demonstrate cooperative targeting within a site spanning nucleotides 158 to 189 of the NRG1 3’ UTR (Figure 5A). NRG1 has a highly conserved 3′UTR sequence overall, with the target sites for the miRNAs showing high sequence conservation, particularly within the seed site, suggesting strong potential for targeting a broad range of hosts using these sites. Closer examination of the sequence conservation reveals that not only the seed regions are conserved across potential hosts, but the deep conservation extends towards the 3′ end of the miRNAs.

**Figure 5.**
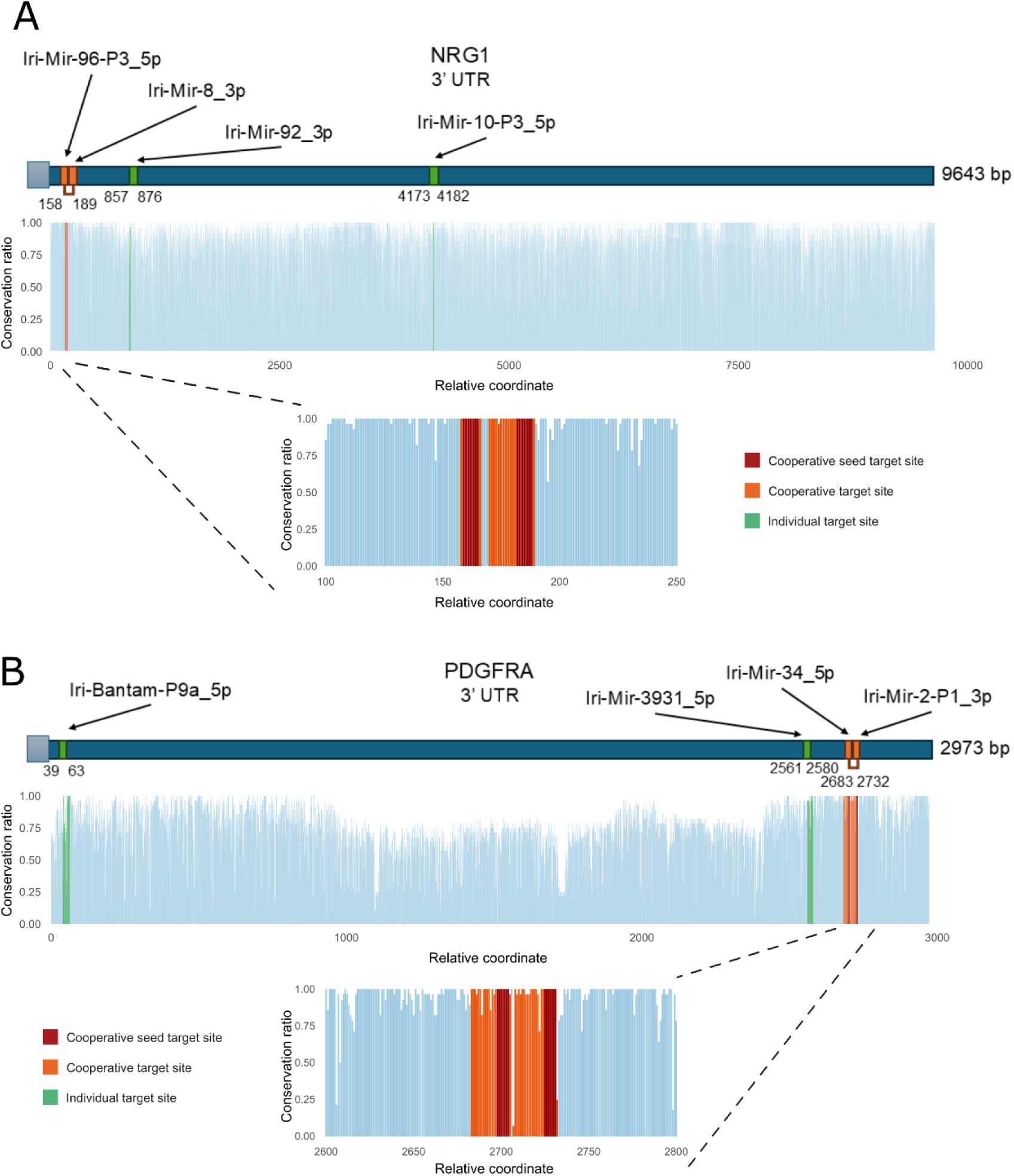
Cooperative effect of core Ixodes ricinus salivary microRNAs on NRG1 and PDGFRA. **A)** Target sites of core I. ricinus salivary miRNAs in the NRG1 gene. Seed target sites of cooperating miRNAs are shown in red, cooperative target sites in orange, and individual target sites in green. Below the schematic representation of the 3′UTR, a bar plot illustrates the conservation ratio (Y-axis) at each position of the 3′UTR (X-axis). A magnified view of the region containing the cooperative target sites is also provided. **B)** Target sites of core I. ricinus salivary miRNAs in the PDGFRA gene. For a detailed description of the visual elements, refer to Figure 5A.

Similarly, PDGFRA is targeted by four tick miRNAs: Iri-Bantam-P9a_5p, Iri-Mir-3931_5p, Iri-Mir-34_5p, and Iri-Mir-2-P1_3p. Among these, Iri-Mir-34_5p and Iri-Mir-2-P1_3p exhibit cooperative binding in a site located between nucleotides 2683 and 2732 of the PDGFRA 3’ UTR (Figure 5B). The cooperative target sites are located within a segment of the 3′UTR that exhibits a particularly high degree of sequence conservation, which suggest their potential utility for broad targeting of multiple host species.

Interestingly, according to KEGG database (Kanehisa et al., 2025), both NRG1 and PDGFRA are involved in the MAPK and PI3K-AKT signaling pathways (Figure 6). Moreover, multiple additional genes within these pathways also possess predicted and conserved target sites for tick miRNAs, suggesting that the tick may synergistically disrupt these host defense mechanisms through coordinated miRNA activity.

**Figure 6.**
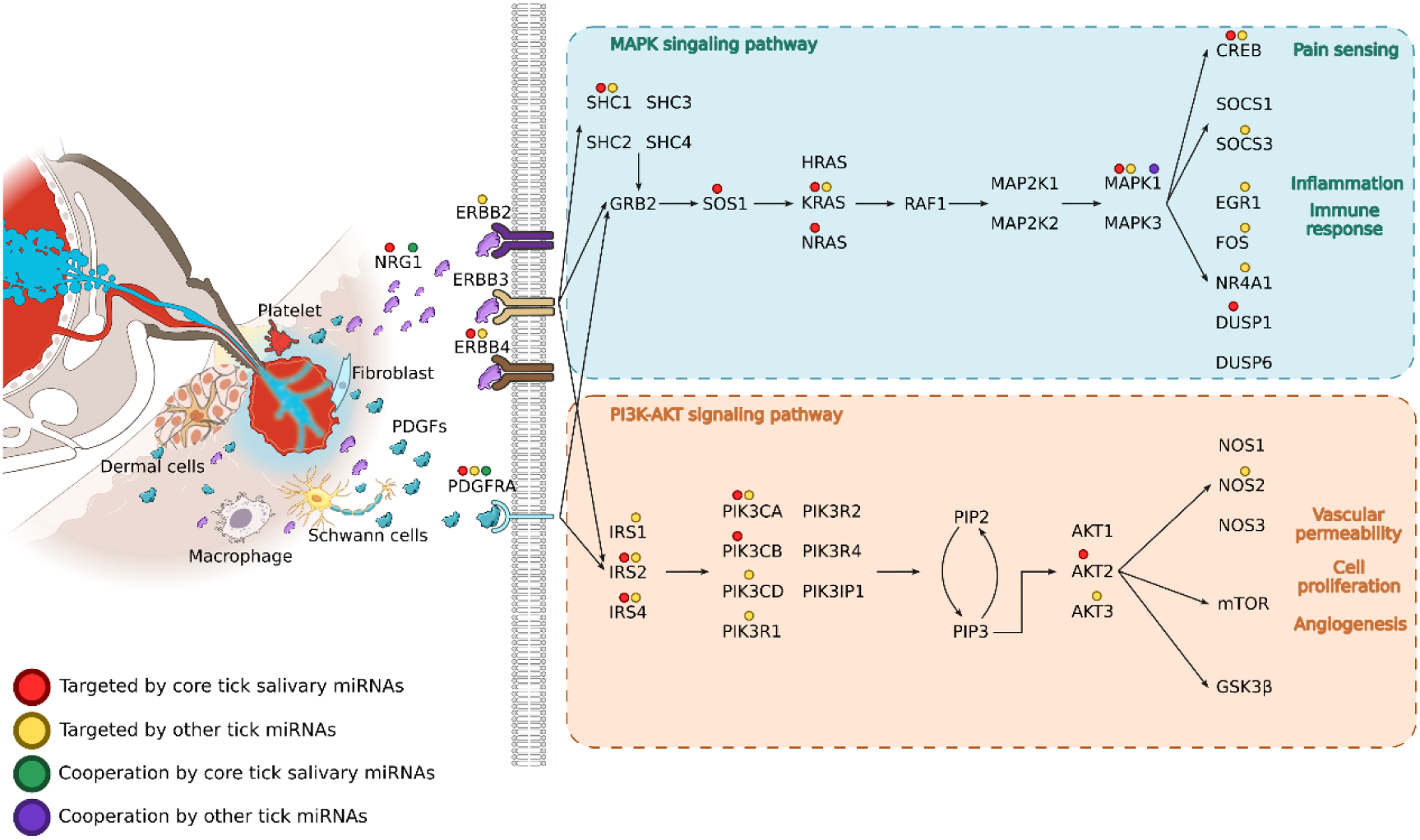
Schematic representation of the roles of NRG1 and PDGFRA at the tick–host interface. This illustration depicts how NRG1 and PDGFRA might modulate the MAPK and PI3K-AKT signaling pathways, highlighting their involvement in the regulation of host defense mechanisms against the tick. Colored circles indicate the presence of individual or cooperative miRNA target sites by core salivary and other I. ricinus miRNAs across the genes shown in the diagram.

## Discussion

The importance of tick saliva proteins in the feeding process has been studied for nearly 4 decades. Tick salivary proteins are not manipulating host gene expression programs but affect different aspects of the host homeostatic response to tick feeding such as wound healing of the host epidermis or host immunity against tick salivary constituents (Kleissl et al., 2025). Nevertheless, the host raises humoral and cellular immune responses against a feeding tick and its tick salivary proteins (Doehl et al., 2025). This is not the case for tick salivary microRNAs, which are non-proteinaceous and non-antigenic. Thus, although they are present in tick saliva, they typically evade immune surveillance. This characteristic, together with their ability to directly manipulate host gene expression programs, makes them interesting candidates for mediating sustained host modulatory effects. Such sustained effects may arise because (i) tick salivary microRNAs “fly under the radar” of the host immune system, and (ii) they can directly modulate gene expression — a property not observed for tick salivary proteins to date. Little is known about the effects of tick microRNAs in the vertebrate host cell homeostasis, especially as far as it concerns the manipulation of important host gene expression programs (Hackenberg & Kotsyfakis, 2018; Nawaz, Malik, Zhang, Hassan, et al., 2020). One of the reasons for this lack of information is the fact that for a large-scale, genome-wide detection of tick microRNA targets in the host transcriptome we rely on computational predictions which however suffer from a high number of false positives (Fridrich et al., 2019; Pinzón et al., 2017). Herein, by applying a conservation filter to the target predictions, we observe: (i) a statistically significant difference between tick microRNA targets (11,746) and their negative control sequences (7,454) and (ii) 12 tick microRNAs that have an outstanding ratio of conserved targets, i.e. a much higher number conserved targets in the host transcriptome, i.e. a much higher number of conserved targets than expected by chance alone. Conservation of the target sites for these 12 tick microRNAs in a wide array of host species is important for the completion of tick lifecycle because it is known that ticks parasitize different hosts depending on their developmental stage and host availability.

We additionally demonstrate that 11 out of these 12 microRNAs are encoded by very ancient genes, present in all bilaterian animals. This means that the specific 11 microRNAs not only they are present in both ticks and all possible host species, but they have identical seed sequences in both ticks and the hosts they parasitize. The consequence of this observation is that mimicry of host microRNAs (the ability to resemble or functionally substitute a host miRNA (Bader et al., 2011; Thorsen et al., 2012)) might play an important role in microRNA-mediated manipulation of host gene expression. Our data propose that tick microRNAs compete with host microRNAs for binding in host mRNA sites which are under selective pressure for microRNA efficiency in the host. This mimicry offers an explanation about how even a low concentration of tick microRNAs can have an impact on gene regulatory programs in the host. Collectively our finding suggests that the broad host range of most tick species may trace back to early Parasitiform evolution, when tick microRNAs—identical in most cases to those in extant species—first engaged target sites in functional regions of their hosts. Functional sites are generally under negative selection, assuring conservation and their presence in all descendent species. Modern ticks in this way would have benefited from the strong diversification of mammals, all with the inherited conserved target regions counting thus with a molecular framework enabling exploitation of a wide diversity of hosts.

Our previous work showed that these microRNAs are present in tick saliva (Hackenberg et al., 2017) and herein we show that their predicted targets are not a random selection but are significantly enriched in several relevant pathways related to host homeostasis such as wound healing, inflammation, pain perception and neurotransmission. When we further enforced cooperativity in the action of these tick microRNAs with respect to their binding to host mRNA targets—an interaction known to enhance repression strength (Briskin et al., 2020; Sætrom et al., 2007)—we predicted an effect of tick microRNAs on 22 host genes expressed in human skin and neural cells, i.e., in cell types present at the tick bite site. At the protein level, two hub genes PDGFRA and NRG1 with a high number of interactions become apparent. Interestingly, the function of these genes converges in the MAPK signaling and PI3K-AKT signaling pathways (Kanehisa et al., 2025) involved in host defense mechanisms, such as pain sensing, or inflammation (MAPK signaling) (Dong et al., 2002; Ji et al., 2009; Moens et al., 2013; Shao et al., 2023) and vascular permeability or angiogenesis (PI3K-AKT) (Karar & Maity, 2011; Marnin et al., 2025; Teng et al., 2021).

Beyond tick–host interactions, the role of microRNAs in cross-species—and even cross-kingdom—gene regulation (e.g., insect–plant or virus–host) has become increasingly evident across diverse biological systems (Arcà et al., 2019; Buck et al., 2014; Cano-Argüelles et al., 2023; Chowdhury et al., 2024; Hackenberg et al., 2017) over the past decade. Parasites can stably transfer microRNAs, protected within membrane-bound extracellular vesicles, into host cells. The central questions remain: which host genes are targeted, and how can a seemingly limited repertoire of parasite microRNAs achieve substantial regulatory effects? Here, we present evidence that tick microRNAs preferentially target conserved regions of host genes—regions of functional importance in the host and therefore protected by negative selection. This rationale provides a strong basis for applying phylogenetic footprinting to minimize false-positive predictions. Moreover, by incorporating cooperativity, we uncover a mechanism that may be particularly critical in parasite–host interactions. Taken together, we are confident that the workflow introduced here, demonstrated for *I. ricinus*–*H. sapiens* interactions, offers a broadly applicable framework for investigating the regulatory role of small RNAs across diverse parasite–host or trans kingdom communication systems.

## Methods

### Data collection

We obtained 3’ UTR regions for canonical transcripts from *Homo sapiens* using BioMart from Ensembl, version 112 (Harrison et al., 2024). The usage of only one transcript per gene, the canonical transcripts avoid redundancies that can arise when using 3’UTRs from different transcripts of the same gene. To obtain the most reliable and updated *I. ricinus* miRNA set, we expanded and refined the previously available *I. ricinus* miRNA complement (GenBank accession numbers MF061606-MF061678). This complement was constructed using an old version of the *I. ricinus* reference genome (Cramaro et al., 2015), the *I. scapularis* genome (Gulia-Nuss et al., 2016), and transcriptomic data from previous studies (Hackenberg et al., 2017). We expanded the old complement by identifying miRNAs in the newly sequenced *I. ricinus* genome (Cerqueira de Araujo et al., 2025) using sRNAbench miRNA annotation protocol (Aparicio-Puerta et al., 2022). The new catalog of *I. ricinus* miRNAs contained 162 mature miRNAs. Of these, 74 could be assigned to *Ixodes scapularis* miRNAs listed in MirGeneDB and were named according to the *I. scapularis* miRNA nomenclature. The remaining 88 were notated as “Novel”.

### Target prediction and conservation

The consensus target prediction involving the algorithms PITA, seed, Miranda and TargetSpy (Enright et al., 2003; Kertesz et al., 2007; Sturm et al., 2010) implemented in sRNAtoolbox^17^ was applied to the human 3’ UTR regions. A consensus target is defined as those predicted by at least 3 out of the 4 algorithms. Next, the consensus target sites are further filtered using the degree of conservation calculated by means of the Vertebrate Multiz Alignment & Conservation (100 Species) provided by UCSC (Nassar et al., 2023). To achieve this, a set of 29 potential host species was curated including: *Homo sapiens, Pan troglodytes, Gorilla gorilla gorilla, Nomascus leucogenys, Macaca mulatta, Macaca fascicularis, Chlorocebus sabaeus, Callithrix jacchus, Spermophilus tridecemlineatus, Mus musculus, Rattus norvegicus, Cavia porcellus, Oryctolagus cuniculus, Bos taurus, Ovis aries, Capra hircus, Equus caballus, Felis catus, Canis lupus familiaris, Mustela putorius furo, Myotis lucifugus, Erinaceus europaeus, Sorex araneus, Monodelphis domestica, Falco cherrug, Falco peregrinus, Ficedula albicollis, Columba livia, Anolis carolinensis*.

Using *H. sapiens* genome as reference, the conservation score was calculated for each position in the target site. The conservation score per position is defined as the number of species with a conserved base (i.e. the same base as the one in the human genome) divided by the number of species with information available in this multiple sequence alignment block. Additionally, we determine the taxonomic level at which the position is conserved. Finally, only target sites with a conservation score greater than 0.9 at all positions in the seed region, showing deeper conservation than the family level (Hominidae in this case), were considered for further analysis.

### Taxon classification

The node of origin of the mature *I. ricinus* microRNA sequences was established applying the seed to define miRNA families. MirGeneDB 3.0 (Clarke et al., 2025), which contains 114 species covering all mayor metazoan groups, was used as reference set. We first extract the seed sequence (position 2-8) of all guide and passenger strand sequences from all MirGeneDB entries and group them together to record all species having this seed sequence. Next, by employing the information contained in the NCBI (Sayers et al., 2025) taxonomy file we determine the last common ancestor for a particular seed sequence (the node of origin) in the following way. For each taxonomic node of the reference species (*I. ricinus*) starting at the species level towards deeper taxonomic nodes we calculate the fraction of MirGeneDB species that belong to the node and having the seed sequence. For example, the first analyzed node is Ixodes as the closest species to *I. ricinus* in MirGeneDB is *I. scapularis*, both in the Ixodes genus. The next taxonomic level with representation in MirGeneDB would be Acari where the red spider mite *Tetranychus urticae* joins. After this, the Arizona bark scorpion *Centruroides sculpturatus* (Arachnida class) and the horseshoe crab *Limulus polyphemus* (*Chelicerata subphylum*) are analysed. The analysis continues towards the deepest metazoan taxonomic node for which microRNAs are reported. The node of origin is defined as the deepest taxonomic node where the fraction is above a threshold of 0.5. If the seed is not in any other MirGeneDB species, or if the fraction is never above 0.5, the seed is defined as *I. ricinus* specific.

### Cooperative effect analysis

Previous studies have shown that a cooperative effect can occur when seed target sites are spaced more than 7 and less than 40 nucleotides apart, with optimal down-regulation observed at distances between 13 and 35 nucleotides (Grimson et al., 2007; Sætrom et al., 2007). This spacing is supported by the bridging function of TNRC6, which facilitates interactions between miRNA-AGO complexes (Briskin et al., 2020). Therefore, we considered seed target sites spaced between 8 and 39 nucleotides apart as potentially cooperative.

### Statistical analysis of target enrichment

miRNAs with a higher number of conserved or cooperative target sites than expected by chance can be interesting starting points for further analysis. In order to assess the statistical significance, we compared *I. ricinus* miRNA target sites with a negative control set of randomized miRNAs. To obtain this set of miRNAs, we used shuffleseq to randomize the sequence of the miRNAs from *I. ricinus*, excluding all random sequences with known seeds in mirGeneDB.

Next, a ranked comparison between *I. ricinus* miRNAs and randomized miRNAs allows us to calculate p-values in the following way: First, we ranked both sets of miRNAs based on the total number of target sites. At each rank position we generate a 2×2 contingency table containing the non-conserved target sites and conserved target sites for tick and random miRNAs. P-values are calculated applying Fisher’s exact test. Correction for multiple testing was applied to the False Discovery Rate (FDR). Odds Ratios are considered as the strength of the biological signal, i.e. how much more conserved target sites a tick miRNA has than expected by chance.

In the case of the cooperative target sites, we ranked both sets by the number of conserved target sites and made the contingency tables using conserved and non-cooperative target sites and conserved and cooperative target sites. As before, Fisher’s test and correction for multiple testing were applied to obtain miRNAs with a higher number of cooperative target sites than expected by chance.

### Functional characterization and protein-protein interaction network

Functional enrichment analysis for tick target genes was made by means of ClusterProfiler (Yu et al., 2012), using pathway and ontology annotations from MSigDB (Liberzon et al., 2015; Subramanian et al., 2005). To ensure comparability with the first functional enrichment analysis in this study, sets of 526 genes with the highest numbers of conserved target sites were selected for functional characterization. In cases where genes had the same number of conserved target sites, selection was performed at random.

The protein-protein interaction network was made using the StringDB (Szklarczyk et al., 2023) module in Cytoscape (Shannon et al., 2003). To analyze a broader picture of the functions of the proteins of interest, we added 50 interactors with a high-confidence threshold of 0.7. Eigenvector centrality of the network was calculated by means of CentiScaPe (Scardoni et al., 2014).

### Analysis of single-cell expression dynamics based on datasets of skin and neural cell types

The genes expression levels across single-cell types from skin tissue and the ratio of single cells expressing these genes were calculated using the single-cell dataset from GTEx (GTEx Consortium, 2013). We calculated the proportion of cells with expression, defined as the median fraction of single cells within each cell type that showed detectable expression for each of the genes in the set. The median expression was computed as the median expression level of all genes in the analyzed gene set for each cell type.

To assess the statistical significance of gene expression patterns, Z-scores and p-values were computed by randomly sampling 1,000 equally sized gene sets, from a background of genes that contained at least one conserved target for the full set of *I. ricinus* miRNAs. The observed expression metrics were then compared to the distribution of metrics expected by chance. P-values and Z-scores were estimated using a Monte Carlo approach.

Single-cell expression data for neural cells were obtained from the Allen Brain Atlas (Allen Institute for Brain Science) (Allen Institute for Brain Science, n.d.). Specifically, we used single-nucleus RNA sequencing data from a dataset comprising over three million cells, sampled from approximately 100 dissections of the adult human brain (Siletti et al., 2023).

## Data availability

The data presented in this study are available within the article and its Supplementary Material. *I. ricinus* miRNAs used in this work are available in Supplementary File 6 (hairpin sequences) and Supplementary File 7 (mature sequences). The *I. ricinus* genome utilized in this study has been published in previous studies (Cerqueira de Araujo et al., 2025) and is accessible at https://bipaa.genouest.org/is/ticks/.

## Author contributions

José María Medina: Formal analysis, Methodology, Visualization, Writing—original draft. Michail Kotsyfakis: Conceptualization, Writing—review & editing. Michael Hackenberg: Conceptualization, Methodology, Formal analysis, Writing—review & editing.

## Acknowledgments

Graphical abstract was created in BioRender. Medina, JM. (2026) https://BioRender.com/5phcyfg.

